# Experimental investigations of the human oesophagus: anisotropic properties of the muscular layer in large deformation

**DOI:** 10.1101/2021.07.18.452813

**Authors:** Ciara Durcan, Mokarram Hossain, Grégory Chagnon, Djordje Perić, Lara Bsiesy, Georges Karam, Édouard Girard

**Author notes:** Email addresses:* (Ciara Durcan), (Mokarram Hossain), (Grégory Chagnon), (Djordje Perić), (Lara Bsiesy), (Georges Karam), (Édouard Girard).

## Abstract

Technological advancements in the field of robotics have led to endoscopic biopsy devices able to extract diseased tissue from between the layers of the gastrointestinal tract. Despite this, the layer-dependent properties of these tissues have yet to be mechanically characterised using human tissue. In this study, the ex vivo mechanical properties of the passive muscularis propia layer of the human oesophagus were extensively investigated. For this, a series of uniaxial tensile tests were conducted. The results displayed hyperelastic behaviour, while the differences between loading the tissue in both the longitudinal and circumferential directions showcased its anisotropy. The anisotropy of the muscular layer was present at different strain rates, with the longitudinal direction being consistently stiffer than the circumferential one. The circumferential direction was found to have little strain-rate dependency, while the longitudinal direction results suggest pronounced strain-rate-dependent behaviour. The repeated trials showed larger variation in terms of stress for a given strain in the longitudinal direction compared to the circumferential direction. The possible causes of variation between trials are discussed, and the experimental findings are linked to the histological analysis which was carried out via various staining methods. Finally, the direction-dependent experimental data was simulated using an anisotropic, hyperelastic model.

## 1. Introduction

The primary function of the oesophagus is to mechanically propel swallowed food, in the form of a fluid bolus, from the pharynx into the stomach through a process called peristalsis [1]. This role can be disrupted by a range of conditions affecting the motor function of the organ, including clinical diseases. Mechanically, 5 peristalsis consists of a combination of passive distentions and active contractions of the oesophageal wall, as well as the interactions of these with the hydrodynamic fluid bolus [2]. Diseases such as diabetes have been found to cause remodelling of the tissue in rats, resulting in an increase in the passive stiffness of the oesophageal wall [3]. To successfully investigate the oesophageal pathophysiology that affects the mechanical function, establishment of the passive mechanical properties of the healthy human oesophagus is first essential [4]. Knowledge of this would also work to enhance the design of medical devices, such as oesophageal stents and endoscopy devices [5, 6], that interact with the passive oesophagus. In addition, recent development in endoscopic biopsy devices in the field of robotics highlights the requirement of quantifying the tissue’s layer-dependent properties [7]. These biopsy devices are designed to extract diseased tissue samples from the wall of gastrointestinal (GI) tract [8, 9], with one such device utilising the fine-needle capillary biopsy technique to pierce through only the inner layer of the wall and retrieve samples from suspected submucosal tumours [10]. The charactisation of the layer-dependent properties of the human GI tract will aid in understanding how the different layers react to an applied stress, and further the development of this minimally invasive diagnostic technique. Further to this, quantification of the passive, layer-dependent properties of the human oesophagus can be used to improve surgical simulations, and also in tissue engineering to cross-reference the mechanical properties of grown tissue against the original [11, 12]. There are a variety of in vivo experimental techniques, such as impedance planimetry and distension inducing probes, however unlike ex vivo experimentation, these techniques do not allow for large deformations and limit the experiments to relatively low strain rates [13, 14, 15, 16, 17, 18].

The vast majority of mechanical experimentation on the oesophagus has been carried out ex vivo on animal tissue [12, 19, 20, 21, 22] due to its wider availability for mechanical testing and reduced ethical issues when compared to human tissue. For instance, Sommer et al. [12] investigated the layer-dependent properties of ovine oesophageal tissue using a series of uniaxial tensile, biaxial tensile, and extension-inflation tests. The tissue exhibited heterogeneous and anisotropic behaviour, and different mechanical properties within the individual layers (the mucosa-submucosa and the muscular layer). The rupture strength of the muscular layer was found to be much lower than that of the mucosa-submucosa layer. Similar to Stravropoulou et al. [19] who also found the muscular layer of porcine oesophagi to be less stiff than the the mucosa-submucosa layer; associating this with the higher collagen content of the mucosa layer. Yang et al. [20] investigated the properties of rat oesophagi and kept the layers intact. In terms of anisotropy, they found the longitudinal direction to be significantly stiffer than the circumferential direction. This is in line with the anisotropic properties established by Sommer et al. [12] and Stravropoulou et al. [19].

The main proportion of experiments carried out to mechanically characterise the human oesophagus have been conducted in vivo using techniques that measure the distensibility of the tissue [13, 14, 15] or using ultrasonic probes [16, 17, 18]. For instance, Orvar et al. [23] found that the circumferential wall tension of the oesophagus increased exponentially with intraluminal pressure. The same observation was obtained also by others in the field [13, 14, 17, 18]. Patel and Rao [14] established the non-uniform distribution of stress that appears along the oesophagus, highlighting the importance of investigating regional discrepancies in regard to its mechanical behaviour. Other studies agree with the Patel and Rao’s findings [18, 24].

Currently, there is very little data on the ex vivo mechanical behaviour of the human oesophagus [24, 25]. Experiments on animal tissue are plentiful, and while animal tissue may provide a good representation of how human tissues might behave, this data cannot be used to accurately model human tissue and successfully enhance applications in medicine. It is therefore essential to obtain experimental data from human oesophageal tissue for this purpose. Egorov et al. [25] conducted a series of experiments on the human gastrointestinal tract, including the oesophagus. They experimented on fresh human cadavers, tested within 24 hours after death. The tissues were stored in a physiological saline solution at 4°C prior to testing and their samples were preconditioned. They studied only the distal third of the oesophagus with the layers intact, and tested only in the longitudinal direction. Investigation into the human oesophagus was limited due to the large collection of tissues being tested. The main conclusions obtained were that the tissue, with layers intact, was found to have a high stressibility and exerted a maximal stress of 1200 kPa. Note that Egorov and co-workers [25] did not consider the layer-dependent properties of the human oesophagus and only investigated the behaviour of one direction. Furthermore, Vanags et al. [24] investigated the effect of pathology and ageing on the mechanical properties within different regions of the oesophagus, comparing the results to healthy tissue. The fresh oesophagi, which were studied with the layers intact, were cut into rectangular specimens in both the circumferential and longitudinal directions and subjected to uniaxial tension until rupture. They found that all oesophagi displayed anisotropic behaviour, with higher resistance in the longitudinal direction. With age, the modulus of elasticity of the tissue wall was found to increase. The cervical part of the oesophagus displayed the highest ultimate stress and strain when compared with the other two regions (thoracic and abdominal). Note that Vanags and co-workers [24] investigated both directions of loading, but did not consider the layer-dependent properties of the tissue.

There is a lack of existing experimental studies into the layer-dependent properties of the human oesophagus regarding its mechanical behaviour. This paper aims to give new experimental data on oesophageal tissue by investigating the properties of the muscular layer of the human oesophagus through a series of uniaxial tensile tests, performed in both the longitudinal and circumferential directions. In the first part, the materials and experimental methods are outlined, including the histological analysis. Next, the experimental and histological findings are presented. Finally, the results of the experiments are discussed by means of anisotropic, hyperelastic modelling.

## 2. Experimental methods

### 2.1. Anatomy and histology of the oesophagus from literature

The oesophagus is an organ of the digestive system located in the the thoracic cavity, as seen in Figure 1a. It is a primarily mechanical organ consisting of several distinct histological layers, as seen in Figure 1b. The mucosa is the inner most layer and consists of three separate layers not visible on the diagram: the epithelium, a non-keratinized stratified squamous covering the lumen of the oesophagus, the lamina propria, a thin layer of connective tissue, and the lamina muscularis mucosae which is a thin layer of muscle tissue [26]. The submucosa is a layer of dense, irregular connective tissue made up of elastin and collagen fibres, containing veins, lymphatics and the submucosal plexus. The muscularis propia layer consists of muscular fibres arranged longitudinally and circularly. The longitudinal muscle fibres are within a more superficial layer, next to the circular fibres which are situated more deeply, as seen in Figure 1b. The longitudinal muscle fibres are gathered laterally in the superior portion of the oesophagus; however, they expand and surround all surfaces as one moves inferiorly down the oesophagus, becoming the strongest in the inferior third of the tissue [26]. Within the superior third of the oesophagus, the circular muscle fibres are elliptical in shape, becoming more circular as one moves inferiorly down the tissue. As well as the orientation, the type of muscle fibres also changes along the length of the tissue. The muscle tissue within the first quarter of the oesophagus is striated, with a mix of striated and smooth muscle fibres within the second quarter. The inferior half of the oesophagus consists of smooth muscle only. A thin layer of connective tissue exists between the two muscular layers (the circular/longitudinal junction) containing the vast proportion of collagen fibres present in this layer [12]. The most superficial layer of the oesophagus is the adventitia, which is formed of loose connective tissue and supports the organ’s position in the thorax [26].

**Figure 1.**
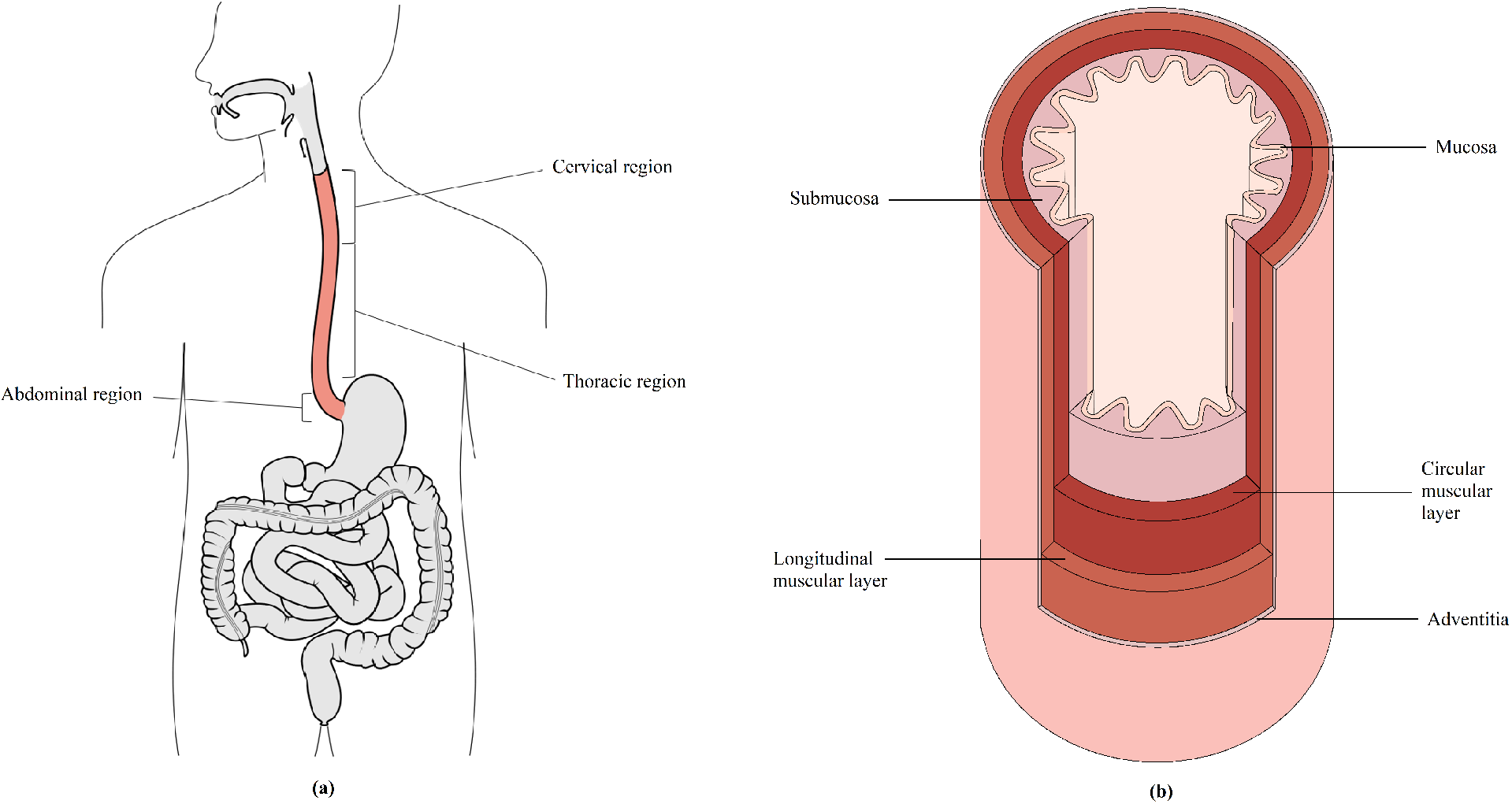
Diagram of the oesophagus showing its position in relation to the rest of the body (**a**) [28] and a segment showing its histological layers (**b**).

The oesophagus is also made up of several distinct regions. These include the cervical, thoracic and abdominal regions, as seen in Figure 1a. The cervical region is the superior region and is 5 - 6cm in length. At its narrowest point, the cervical region has a luminal diameter of 1.4 - 1.5cm, with the remaining regions of the tissue having a luminal diameter of approximately 2cm. Below the cervical region is the thoracic region, which accounts for the majority of the oesophagus, and is 16 - 18cm in length. The inferior and smallest region of the tissue is the abdominal region which is 1 - 2.5cm in length, depending on the size of the person [26].

Once explanted, the oesophagus can be easily separated into two layers; a muscularis propia layer and a mucosa-submucosa layer, allowing for separate mechanical characterisation of the tissue layers. The following findings are from Stavropoulou et al. [27] who conducted histological observations on rabbit oesophagi. The collagen content in the muscle layer was found to be 15.02% in the longitudinal direction and 9.54% in the circumferential direction. The collagen content for the mucosa-submucosa layer in the longitudinal and circumferential directions were 20.25% and 18.17%, respectively. The elastin content in the muscle layer was 3.48% in the longitudinal direction and 1.16% in the circumferential direction. The elastin content for the mucosa-submucosa layer in the longitudinal and circumferential directions were 10.45% and 5.43%, respectively. They found both the collagen and elastin content to be significantly lower in the muscular layer than the mucosa-submucosa layer in both directions. For both layers, elastin content was significantly higher in the longitudinal direction than in the circumferential one. There was also more collagen content in the longitudinal direction than the circumferential direction for both layers, however the difference was not significant.

### 2.2. Sample extraction

The whole human oesophagus was extracted by means of dissection at the Laboratoire d’Anatomie Des Alpes Françaises, Grenoble, France. The oesophagus was retrieved from an embalmed cadaver due to restrictions caused by the COVID-19 pandemic, wherein fresh cadavers were not available for dissection. Cadavers were embalmed with a formalin solution (ARTHYL) injected into the carotid artery and drained from the jugular vein, and then preserved in a 4°C refrigerated room. All cadavers were required to present a negative COVID test before being allowed for dissection.

A median phreno-laparotomy was performed up to the umbilicus, as well as a left cervical approach following the edge of the sterno-cleido-mastoid muscle. Starting from the stomach, the abdominal part of oesophagus was individualised from the hiatus in the diaphragm. The left triangular ligament, which connects the diaphragm and the posterior surface of the left lobe, was freed, allowing the liver to be reclined. The small omentum was then sectioned off so that the stomach could be freed and hooked up to locate the abdominal oesophagus. The visceral peritoneum in front of the oesophagus was dissected, then a phrenotomy was performed. Dissection of the oesophagus continued in a cranial direction until the pulmonary hilum with its triangular ligament was severed. A right thoracic approach was chosen for the rest of the dissection, allowing the oesophagus to be individualised without being obstructed by the aorta and the heart in the left part of the mediastinum. The right lung was then redirected to the front and the pulmonary hilum was above the oesophagus. The great azygos vein was also dissected on the right edge of the oesophagus. The abdominal oesophagus was sectioned by making an incision in the fundus of the stomach. The trachea was then sectioned in front of the oesophagus due to it preventing access to the cervical part of the oesophagus. Finally, the cervical oesophagus was sectioned below the pharynx and the whole oesophagus extracted.

This study was performed in compliance with French regulations on postmortem testing, and the protocol approved by a local scientific committee from Grenoble-Alpes University.

### 2.3. Histology

Samples for histological analysis were obtained in the transversal and longitudinal planes of the entire tissue, and the coronal plane of muscularis propria layer. Samples were conserved in formaldehyde, fixed first in formalin 10% during 24 hours at 4°C and then embedded in paraffin according to usual protocol (Canene-Adams, 2013). Sections of 3μm were then realised with a microtome Leica RM 2245 (Wetzlar, Germany). The slides were then stained with Hematoxylin Eosin Saffron (HES) to see the nucleic acids and connective tissue (amongst other collagen), with Orcein staining to highlight elastin fibres or with Sirius Red to highlight all types of collagen and muscular fibres.

### 2.4. Sample preparation

The oesophagus was approximately 27cm in size, as seen in Figure 2. In preparation for the testing, the oesophagus was cut into its three separate regions (cervical, thoracic, and abdominal), highlighted in Figure 2, by cutting along the circumferential direction. The thoracic region was then cleaned by removing any excess connective tissue. At this point, some samples were removed for histological analysis, as described in Section 2.3. The remaining tissue was then used for mechanical testing.

**Figure 2.**
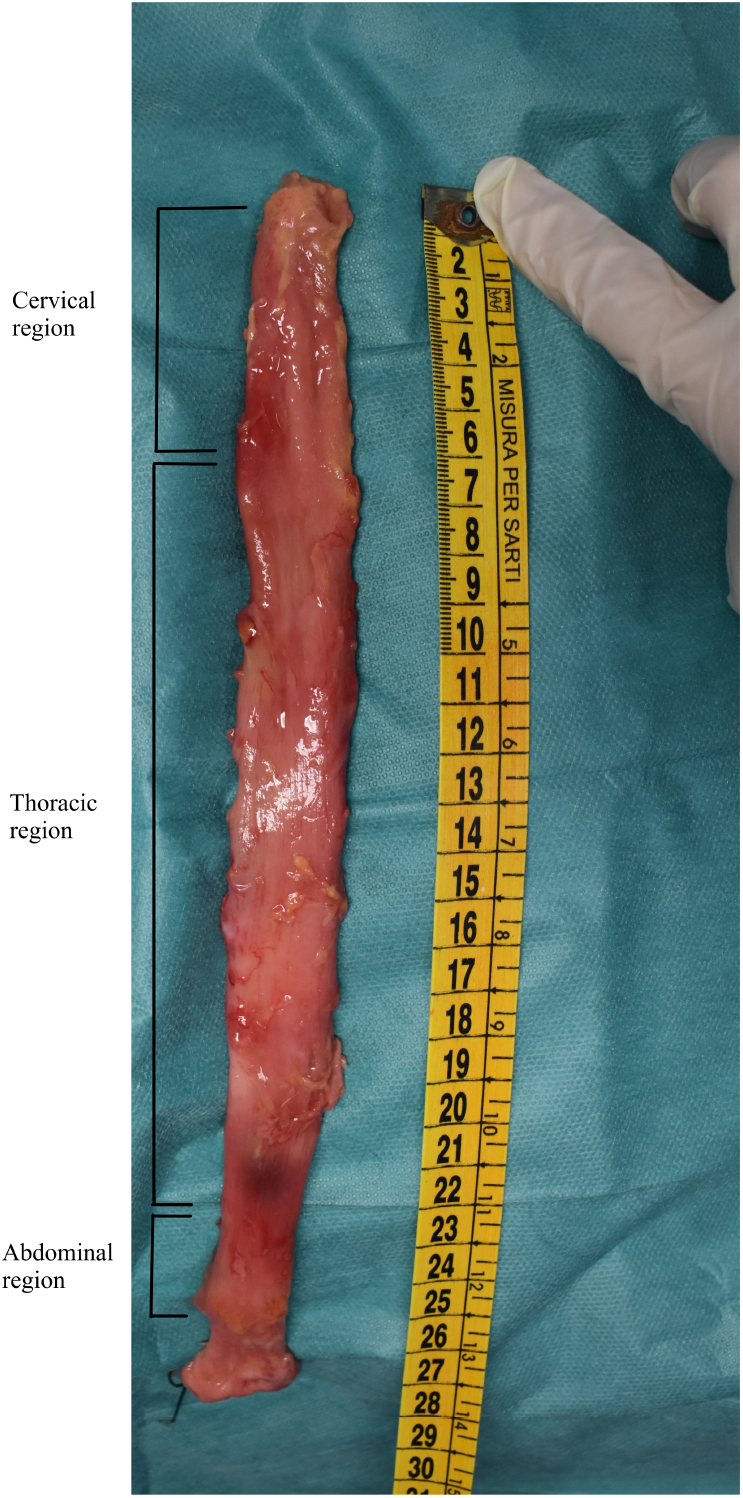
An entire human oesophagus post explantation with its regions labelled.

The distinct layers of the tissue can be seen in Figure 3a. The separation of layers was initiated by cutting carefully through the outer, muscular layer along its longitudinal length. This opening was used to separate the muscular layer from the mucosa-submucosa layer through a series of small cuts to the loose connective tissue binding the layers together, as seen in Figure 3b. The muscle layer was then unravelled and rectangular specimens approximately 20.00mm x 3.80mm (length x width) in size were cut in both the longitudinal and circumferential directions, as seen in Figure 4a, from the inferior end of the thoracic region where the muscle fibres are more evenly distributed. In spite of this, natural variations in the layer were still prevelant, causing difficulty when attempting to cut as consistent samples as possible. Time was taken to cut the samples due to the soft and delicate nature of the tissue, with specical care taken to cut them as parallel as possible to the orientation of their respective muscle fibres. The testing was completed within 5 days of explantation, during which the tissue was stored in physiological saline solution (0.9% NaCl) in a 4°C refrigerator and new samples cut each day. Before testing, the samples were brought to ambient temperature and were kept moist between tests.

**Figure 3.**
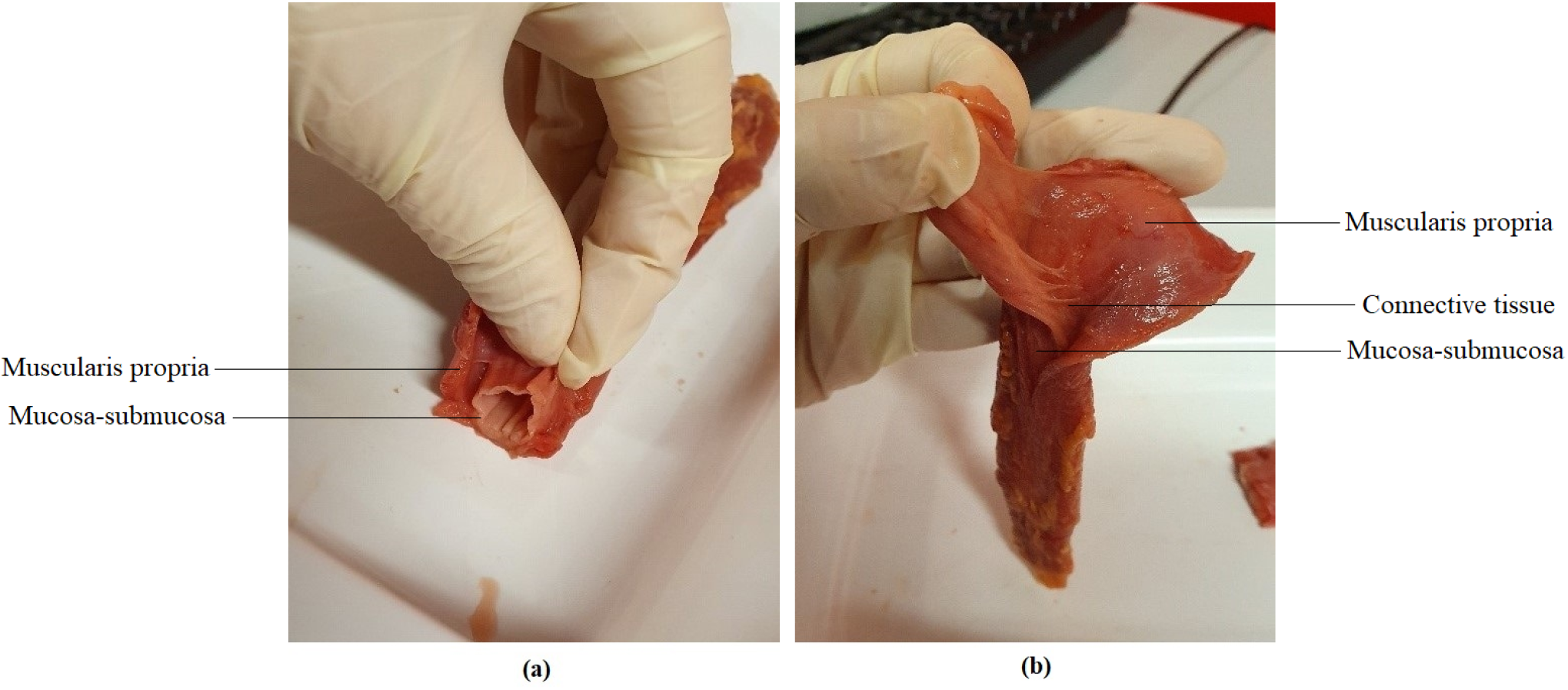
**a** The distinct layers of the oesophagus prior to layer separation. **b** During separation of the oesophageal layers, showing the connective tissue between them.

**Figure 4.**
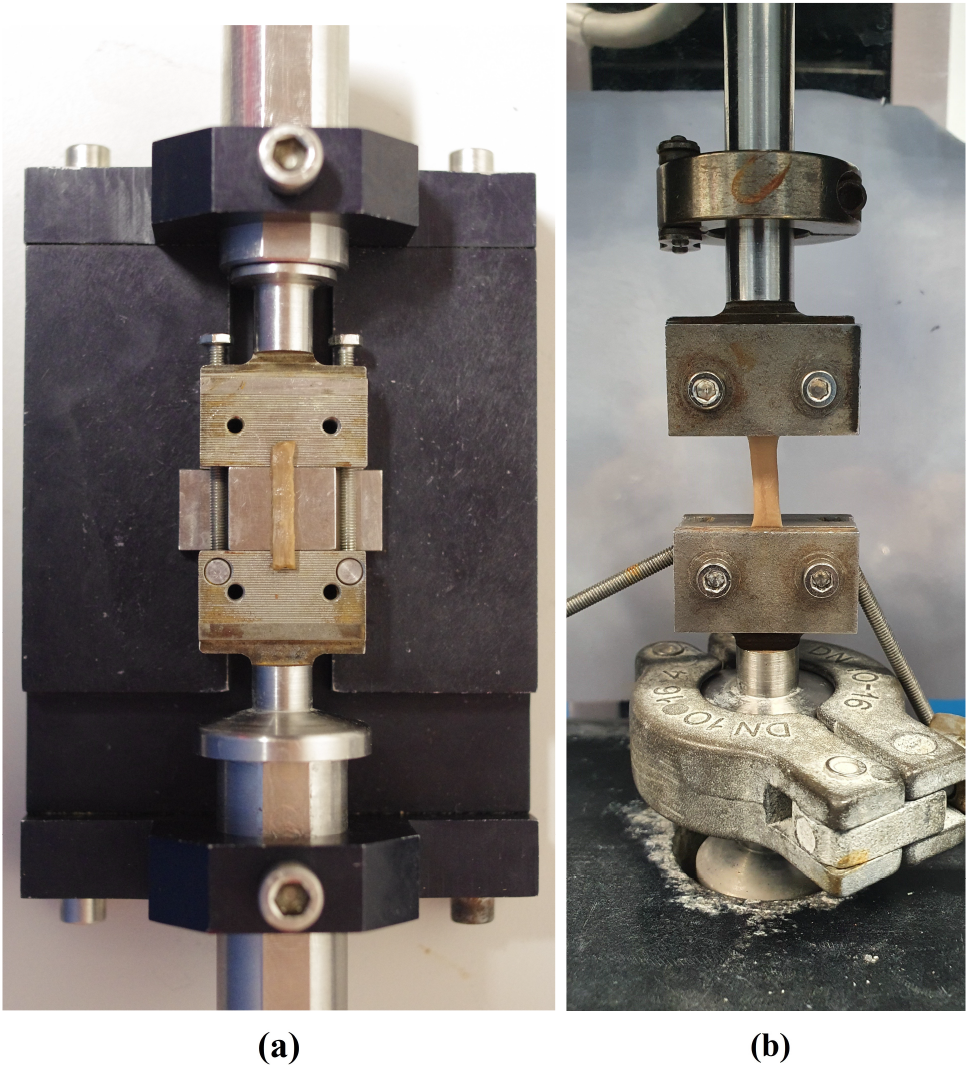
**a** Rectangular sample being loaded between the grips. **b** Sample loaded in the machine.

### 2.5. Experimental setup

The specimens were loaded between the grips using a specially designed device as seen in Figure 4a. A support, 13.9mm in height, was used to keep the soft tissue specimen in place while being secured within the grips. Each sample was placed centrally on the bottom plates while being held in situ by the support. This step took time due to the delicate and moist nature of the tissue causing the sample to stick; proving it difficult to move the sample by the small margins necessary to align it as centrally as possible. The top plates of the grips were then added and the screws tightened partially at first in all four corners to prevent any potential uneven distribution of the sample within in the grips. The screws were then tightened fully to hold the sample securely in place. For consistency, a torque limiter set at 0.5Nm was used to tighten the screws. Long screws, as seen to the right and left of the specimen in Figure 4a, were then tightened to keep the assembly (upper grip, support and lower grip) in place and the specimen from any damage while being loaded into the traction machine. Once setup in the machine, the long screws as well as the support were removed, leaving the specimen loaded within the machine as seen in Figure 4b. At this point, the thickness and width of the specimens were measured using callipers at three separate points and an average was taken. A highly sensitive 25N load cell was fitted to the MTS Criterion model C41 traction machine and used for all tests due to the comparatively low internal stresses of human soft tissues. If the samples buckled while being secured in the grips, the crosshead would be adjusted once the sample was loaded into the machine to reflect its new initial length and the load would be zeroed. The machine and the test parameters were controlled and inputted using the MTS TestSuite user interface software.

### 2.6. Mechanical characterisations

All tests were carried out at ambient temperature and conducted under a uniaxial tensile test condition. The stretch is defined as 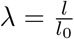, where *l* and *l*_0_ are the current and initial heights of the specimen, respectively. The experimental strain is expressed in terms of stretch, λ, which relates to nominal strain by *ε* = λ – 1. The strain rates are expressed in units of percentage deformation per second (%*s*^−1^).

Similarly, the stress is expressed as the nominal stress (i.e., first Piola-Kirchhoff stress), which is defined as:

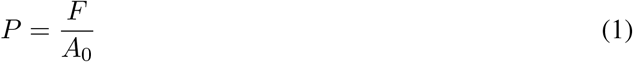

where *F* is the applied force and *A*_0_ is the original, undeformed cross-sectional area. Preconditioning is a common technique used for the mechanical characterisations of non-biological materials such as soft polymers, and is sometimes used in the biomechanical characterisation of soft tissues [29, 30, 31]. However, the technique was decided against in this study due to the applications of the experimental data within areas such as surgical simulations, for which the material response of un-preconditioned tissue is paramount to its function.

Uniaxial tensile tests, wherein the length of the sample is substantially larger than the width, and the force is exerted along a single axis parallel to its length, were conducted until fracture to observe the material response of the tissue. Due to the non-homogeneous distribution and orientation of fibres present in the muscle layer, tests were conducted in both the longitudinal and circumferential directions to observe the effect of direction of loading on the stress response. The tests were also carried out at two different constant strain rates, 1%*s*^−1^ and 10%*s*^−1^, to observe any time-dependent behaviour in each of the directions; a response associated with the viscoelastic behaviour of biological tissues [2]. Each test was repeated at least five times per direction per strain rate to ensure reproducible results.

## 3. Results

### 3.1. Histological analysis of the muscularis propria layer of the human oesophagus

In the transversal plane, the four layers of the oesophagus outlined in Section 2.1 are clearly visible and can be seen in Figure 5a. These layers are also evident in the longitudinal plane, as seen in Figure 5b. Compared to the mucosa-submucosa, the muscularis propria has a lower composition of collagen and elastin fibres. Overall, there are more collagen fibres than elastin fibres in the fibrous composition of the muscularis propria (both circular and longitudinal layers). In the inner circular layer, the collagen fibres form a mesh orthogonal to the axis of the oesophagus. Fibres have a transversal orientation with some oriented along the muscle cells and others towards the lumen. Elastin fibres have a similar orientation in this layer. The fibres are mostly concentrated at the circular/longitudinal junction of the muscularis propria, organised mainly along the axis of the oesophagus. The outer longitudinal layer has its collagen fibres mainly oriented in a longitudinal orientation, with few fibres oriented towards the lumen. The elastin fibres have a longitudinal orientation also, but with a very wavy appearance. The elastin fibres seem to be more abundant in the outer longitudinal layer than in the inner circular layer. The distribution of collagen and elastin fibres within the muscularis propria are summarised in Table 1.

**Figure 5.**
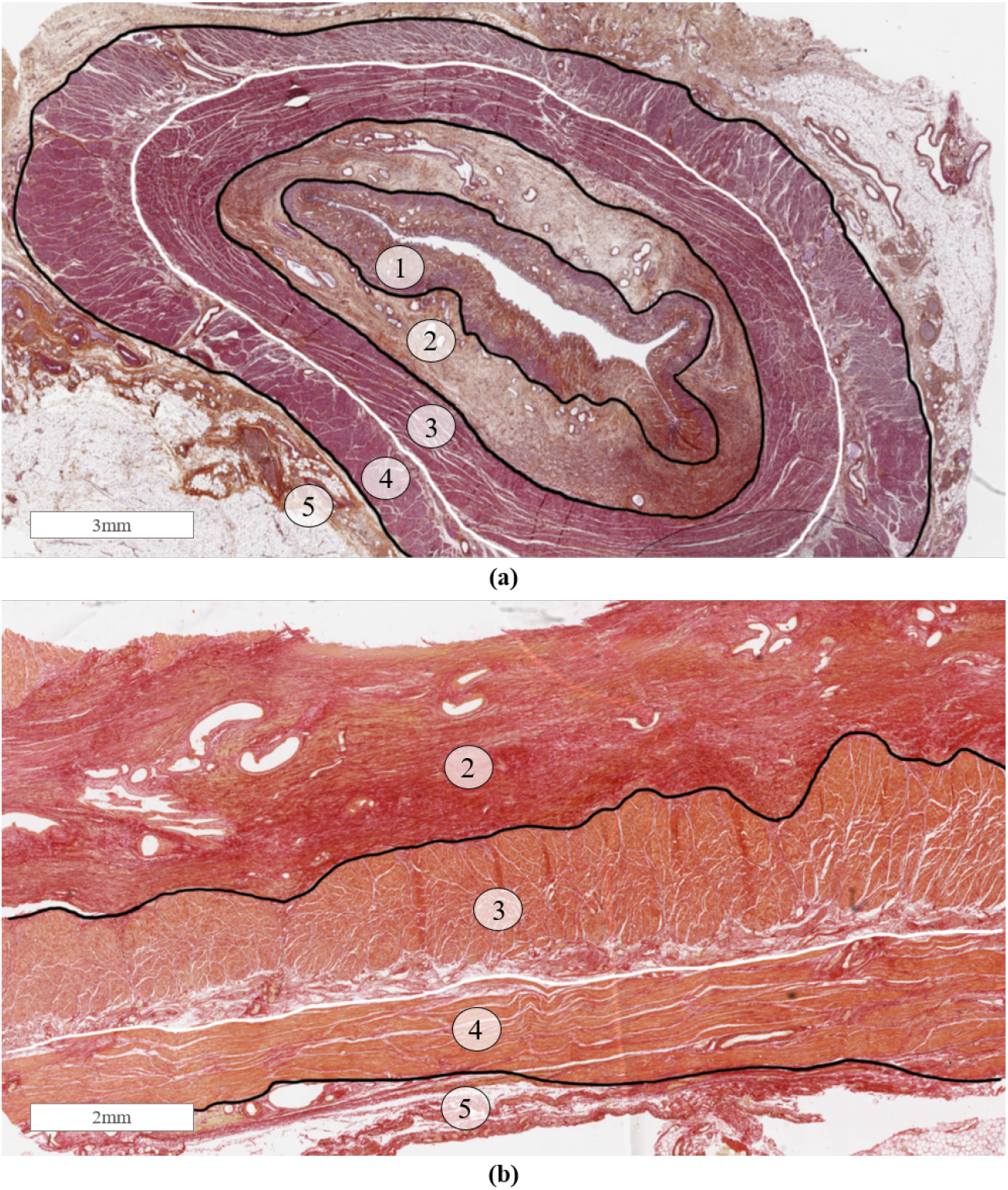
Hematoxylin Eosin Saffron staining in the transversal plane (**a**) and Sirius Red staining in the longitudinal plane (**b**) showing the mucosa (**1**), submucosa (**2**), the circular muscle fibres of the muscularis propria (**3**), the longitudinal muscle fibres of the muscularis propria (**4**) and the adventitia (**5**).

**Table 1:** Distribution of collagen and elastin in the muscularis propria; +, low density; ++++, high density.

| Layer | Collagen | Elastin |
| --- | --- | --- |
| Circular muscle | ++ | + |
| Circular/longitudinal junction | ++++ | +++ |
| Longitudinal muscle | ++ | ++ |

### 3.2. Demographics and variations in experimental samples

The oesophagus was retrieved from an embalmed cadaver stored for 29 days after death while being tested for COVID-19. The patient was male with a total height of 180cm, an age of 73 years and a weight of 90kg. Dimensions such as thickness can vary from sample to sample due to the variable nature of biological tissues. Specimens were obtained from the inferior end of the thoracic region where the muscle fibres are more evenly distributed. However, variations in terms of thickness were still present, as seen in Table 2. The width of the specimens were also seen to vary due to the human error associated with measuring and cutting by hand.

**Table 2:** Mean and standard deviations of the sample dimensions.

| | $1\%s^{-1}$ | | $10\%s^{-1}$ | |
| --- | --- | --- | --- | --- |
|  | Width [mm] | Thickness [mm] | Width [mm] | Thickness [mm] |
| **Longitudinal** | $3.66 \pm 0.19$ | $0.72 \pm 0.09$ | $3.94 \pm 0.28$ | $0.68 \pm 0.05$ |
| **Circumferential** | $3.82 \pm 0.22$ | $0.80 \pm 0.11$ | $3.82 \pm 0.25$ | $0.92 \pm 0.08$ |

### 3.3. Reproducibility in stress-strain data

Variations in results between repeat tests were observed in terms of fracture stretches and stiffness. The fracture stretches of each trial and strain rate can be seen in Table 3. Fracture of the specimen was defined as irreversible structural damage, visible on the stress-strain graph as a sudden reduction in stress. The specimens fractured in various locations, including in the middle of the specimen, at the grip location, or a combination of both. Figure 6 shows the variations in stress-strain data between trials of the 1%*s*^−1^ tests in both the longitudinal and circumferential directions. The variation in stiffness between repeats was found to be larger in the longitudinal direction than the circumferential direction. This is believed to be due to a more uneven distribution of muscle fibres in the longitudinal direction than the circumferential one; wherein the longitudinal muscle fibres are gathered laterally in the superior portion of the tissue, and gradually become more evenly distributed as one moves inferiorly down the tissue. This results in more variations throughout the tissue and therefore the samples. The subsequent graphs are based on the mean average of the three closest curves of the repeated trials for each strain rate and direction.

**Table 3:**
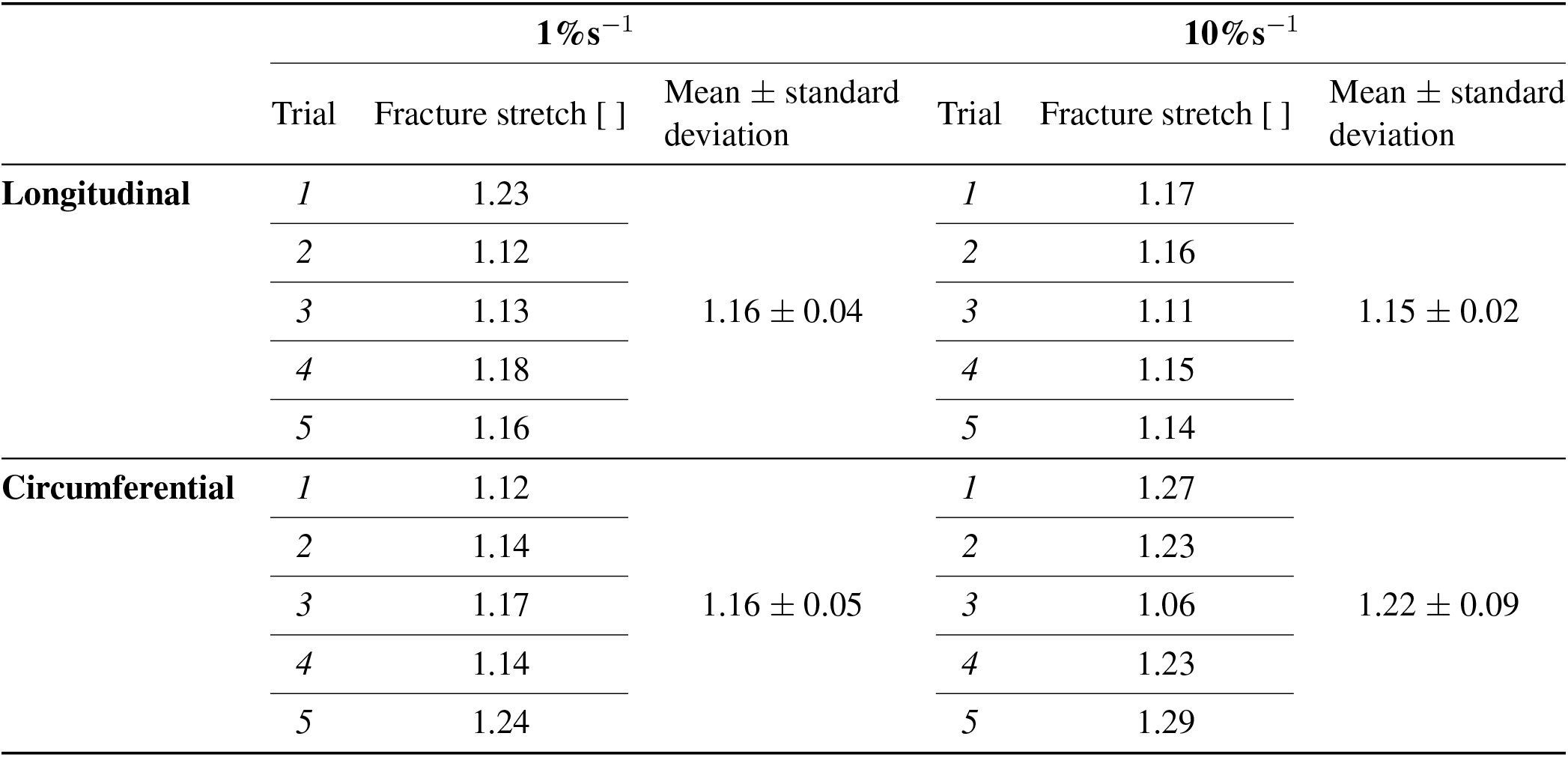
Mean and standard deviations of fracture stretches.

| | $1\%s^{-1}$ | | | $10\%s^{-1}$ | | |
| --- | --- | --- | --- | --- | --- | --- |
| | Trial | Fracture stretch [ ] | Mean ± standard deviation | Trial | Fracture stretch [ ] | Mean ± standard deviation |
| **Longitudinal** | 1 | 1.23 | $1.16 \pm 0.04$ | 1 | 1.17 | $1.15 \pm 0.02$ |
|  | 2 | 1.12 |  | 2 | 1.16 |  |
|  | 3 | 1.13 |  | 3 | 1.11 |  |
|  | 4 | 1.18 |  | 4 | 1.15 |  |
|  | 5 | 1.16 |  | 5 | 1.14 |  |
| **Circumferential** | 1 | 1.12 | $1.16 \pm 0.05$ | 1 | 1.27 | $1.22 \pm 0.09$ |
|  | 2 | 1.14 |  | 2 | 1.23 |  |
|  | 3 | 1.17 |  | 3 | 1.06 |  |
|  | 4 | 1.14 |  | 4 | 1.23 |  |
|  | 5 | 1.24 |  | 5 | 1.29 |  |

**Figure 6.**
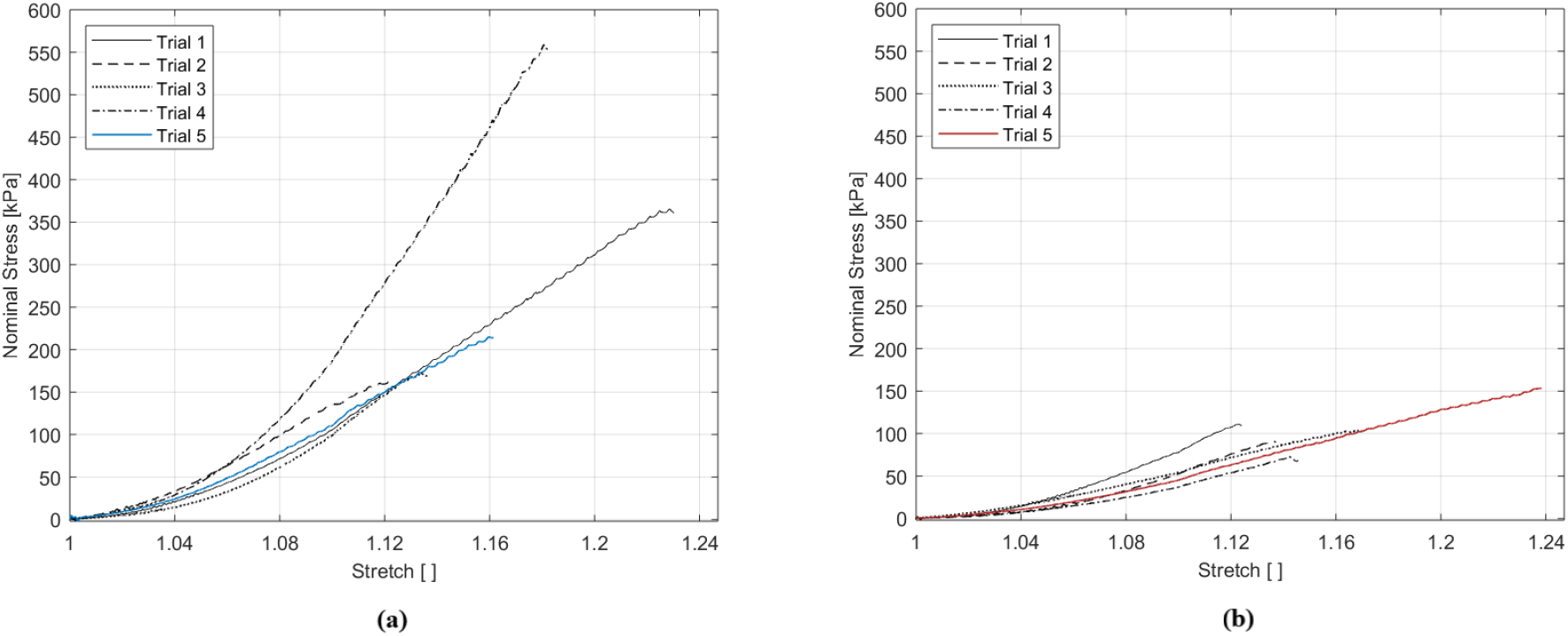
Experimental results of the 1%*s*^−1^ tests showing the variation in the longitudinal direction (**a**) and circumferential direction (**b**).

### 3.4. Anisotropic response

The results comparing the loading directions for the two strain rates are presented in Figure 7. The muscular layer of the human oesophagus displays anisotropic properties at both strain rates. The stress was higher in the longitudinal direction than the circumferential direction for both strains rates; with the stress at 1.1 stretch in the longitudinal direction being approximately double that compared to the circumferential direction for the 1%*s*^−1^ tests, and over 5 times greater for the 10%*s*^−1^ tests.

**Figure 7.**
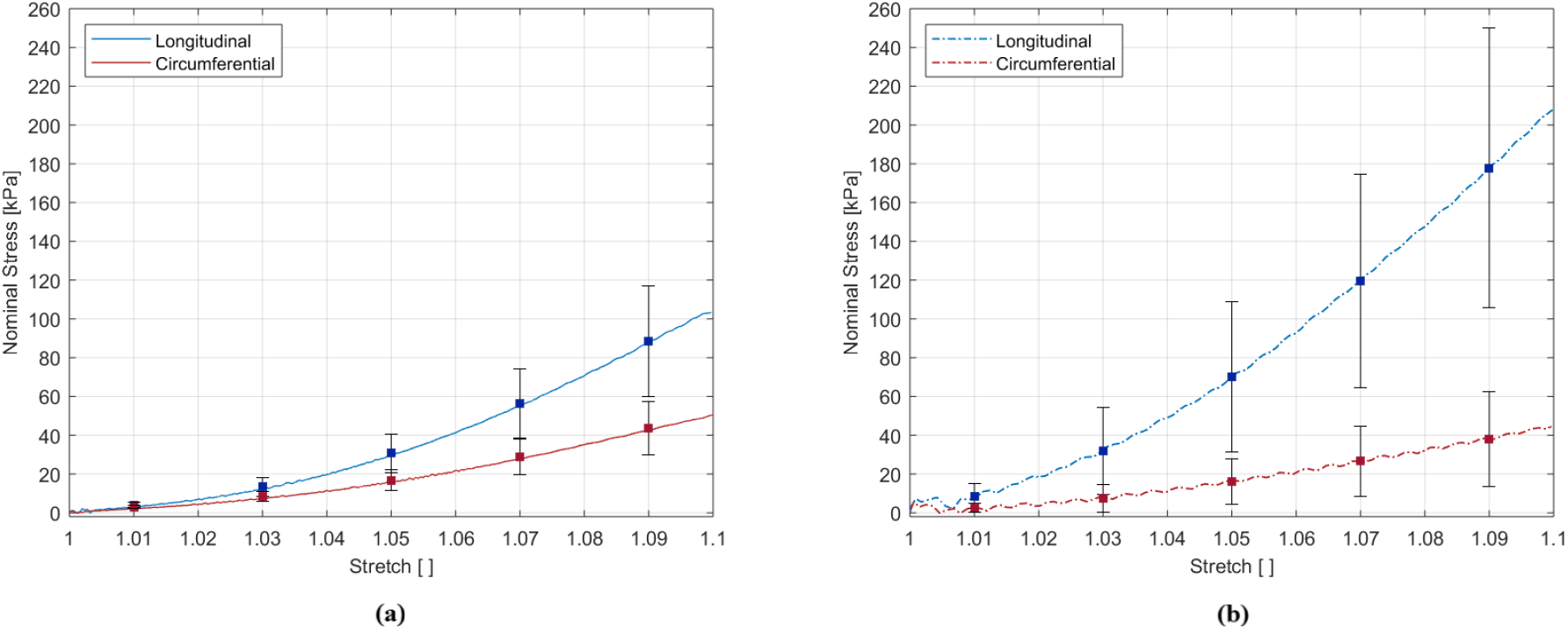
Effect of the direction of loading on the average results at 1%*s*^−1^ (**a**) and 10%*s*^−1^ (**b**).

### 3.5. Effects of strain rate

Figure 8 compares the strain rates for the longitudinal and circumferential directions. The strain rate was found to have little effect in the circumferential direction, while the results in longitudinal direction suggest strain-rate-dependent behaviour. In the longitudinal direction, the stress at 1.1 stretch was approximately double for the 10%*s*^−1^ tests when compared to the 1%*s*^−1^ tests.

**Figure 8.**
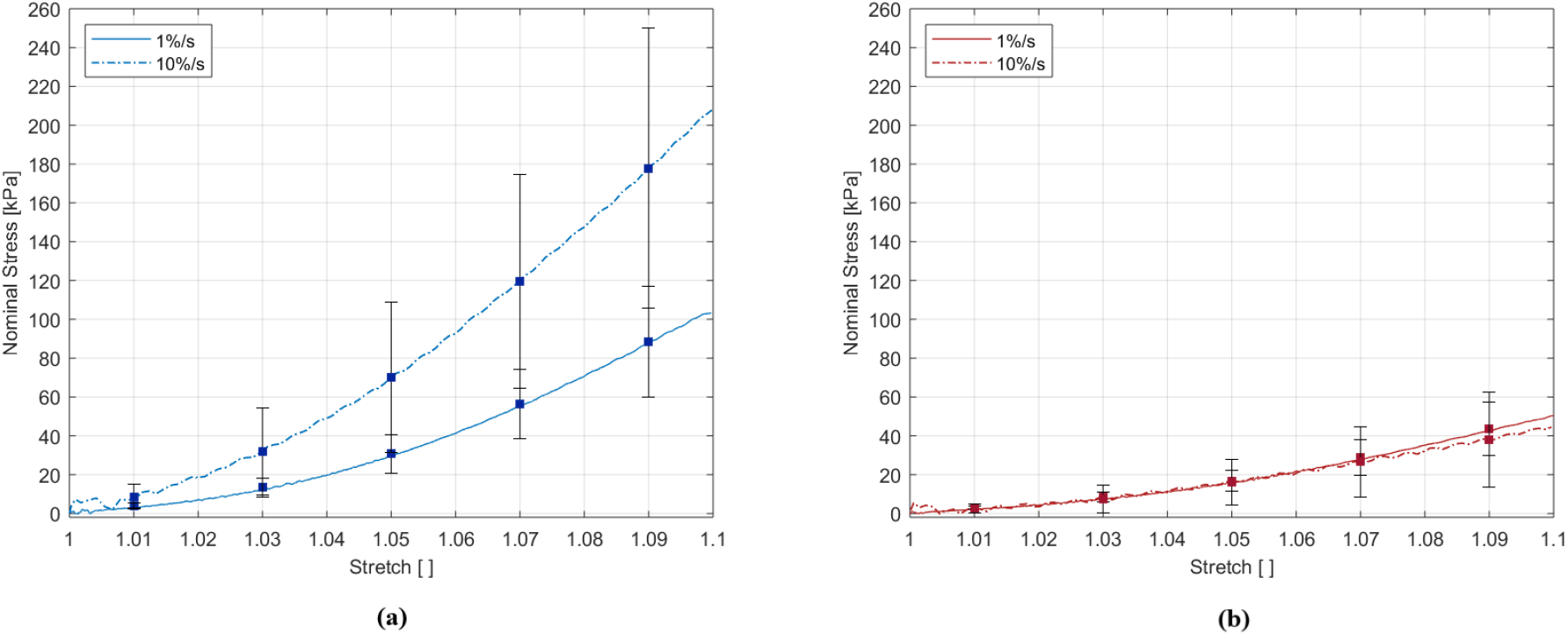
Effect of strain rate on the average results in the longitudinal direction (**a**) and the circumferential direction (**b**).

## 4. Discussion

### 4.1. Hyperelastic description

In this section, we demonstrate the procedure to model the experimentally observed behaviour of the muscular layer of the human oesophagus using an anisotropic, hyperelastic matrix-fibre model proposed for collagen-reinforced soft biological tissues. The hyperelastic behaviour was modelled for both a low strain rate (1%*s*^−1^) and a medium strain rate (10%*s*^−1^) to provide a choice of material parameters for static finite element simulations, depending on the application. The matrix of the tissue has been considered as purely isotropic, with the anisotropy originating from the fibres and their predominant orientations. In this case, the fibres are composed of the collagen networks surrounding and connecting the muscle fibres and, from our histological analysis as outlined in Section 3.1, were observed to sit predominantly in the same direction as their respective muscle fibres. Therefore, the collagen fibres of the muscular layer of the oesophagus are considered to be orientated in the longitudinal and circumferential directions, and are classed as orthogonal. ***N***^(*i*)^ is the direction of each set of the fibres in the undeformed state. For this model, it is assumed that there are two main families of collagen fibres whose effect can be captured by their mean orientations, ***N***^(1)^ and ***N***^(2)^, as seen in Figure 9. For the longitudinal samples, the direction vectors are simply as follows:

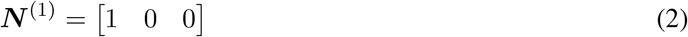

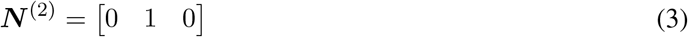

**Figure 9.**
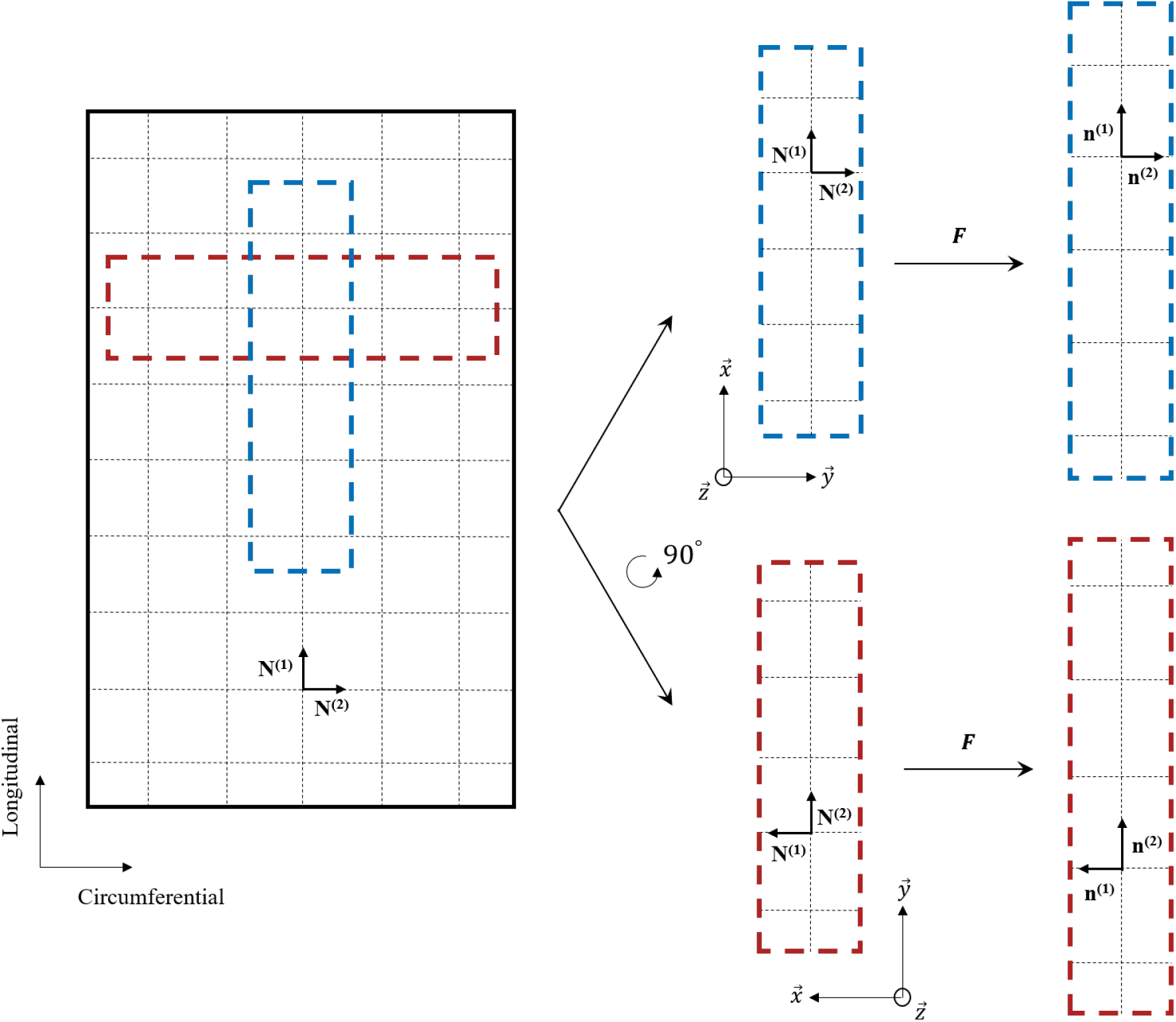
Drawing to illustrate the fibre orientation of the muscularis propria of the human oesophagus based on the histological observations outlined in Section 3.1.

The orientation in the deformed state can be captured at any given time by ***n***^(*i*)^ = ***F N***^(*i*)^, in which ***F*** is the deformation gradient tensor.

A histological based strain-energy function established by Holzapfel et al. [32] is chosen here and defined as:

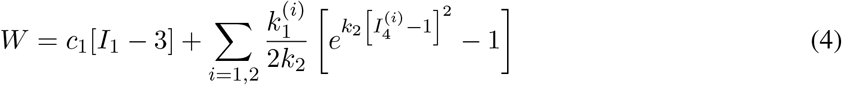

where, *c*_1_ and *k*_1_ > 0 are stress-like material parameters, and *k*_2_ > 0 is a dimensionless parameter. For a comprehensive review on a variety of anisotropic, hyperelastic energy functions, readers are referred to Chagnon et al. [33]. The invariants in Equation (4) are defined as:

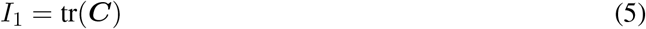

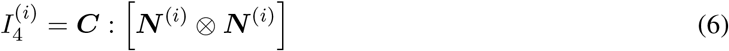

where, ***C*** = ***F^T^ F***. For uniaxial tension, the specimen is loaded in only one direction, i.e. λ*_x_* = λ for the longitudinal samples, while the other two directions are unhindered. In this case, it is assumed that the two families of fibres are active only in tension. Therefore, for an incompressible material and due to the assumption of symmetry, the deformation gradient tensor for uniaxial tension can be written as:

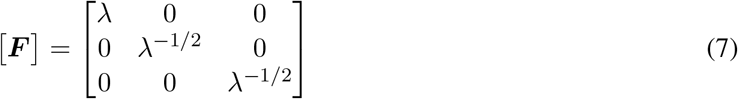

where, λ is the stretch as defined in Section 2.6. In order to find the stress-strain relationship for the case of uniaxial tension, the partial derivatives of the energy function seen in Equation (4) are performed with respect to the strain invariants *I*_1_ and *I*_4_, resulting in the second Piola-Kirchhoff (PK) stress tensor equation for incompressible materials:

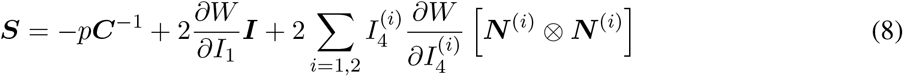

where, *p* is the hydrostatic pressure and ***I*** is the identity tensor, see [34, 35, 36, 37, 38]. The first Piola-Kirchhoff tensor is related to the second Piola-Kirchhoff by:

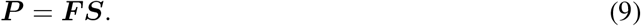

By inserting the definition of hydrostatic pressure for uniaxial tension and the definitions of the invariants *I*_1_, 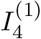 and 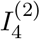, the equations for the one-dimensional first PK stress for each loading direction become:

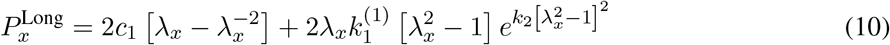

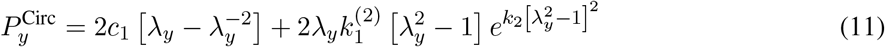

The equation for the first PK stress expressed in terms of λ is now compared with the experimentally observed stress-strain data to identify the unknown parameters *c*_1_, *k*_1_ and *k*_2_.

The experimental data obtained at 1%*s*^−1^ strain rate for the longitudinal and circumferential directions of the muscular layer were fit simultaneously to the one-dimensional forms of the first PK stress from Equations (10) and (11). The calibration results can be seen in Figure 10a, which depict a good fit with the model. Now, the model is validated with another set of data that was not included in the parameter identification process. In this case, the 10%*s*^−1^ longitudinal experimental data was selected. As expected, a higher strain rate yields a stiffer stress value. Hence, for the validation, the already identified parameter *k*_2_ was kept fixed while the stiffness variations were captured by the changing value of the parameters *c*_1_ and *k*_1_, as seen in Figure 10b. The 10%*s*^−1^ circumferential experimental data was then modelled using the *c*_1_ parameter for the higher strain rate. The *c*_1_ parameter represents the stiffness attributed to the isotropic matrix and describes the average stiffness before the activation of the fibres, while the *k*_1_ parameter represents the stiffness of the fibrous anisotropic material components [12]. All parameter values can be found in Table 4.

**Figure 10.**
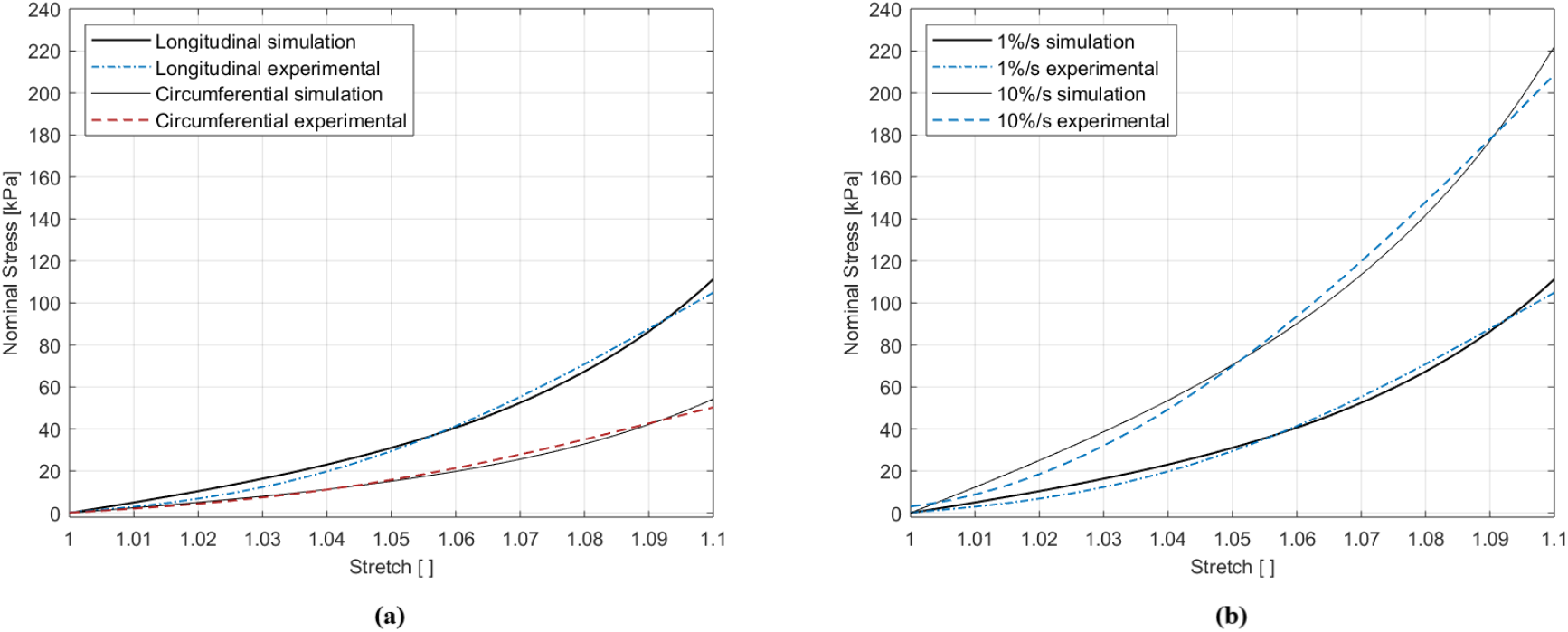
Modelling of the two directions simultaneously at 1%*s*^−1^ (**a**) and modelling of the two strain rates in the longitudinal direction (**b**).

**Table 4:**
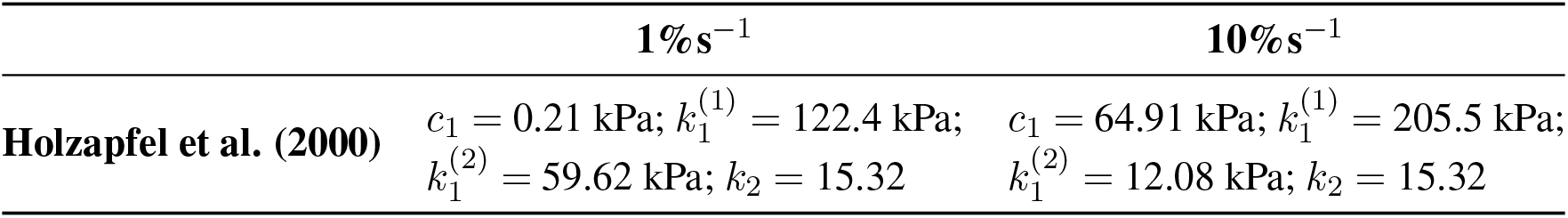
Material parameter values for the longitudinal and circumferential directions at 1%*s*^−1^ and 10%*s*^−1^

| | $1\%s^{-1}$ | $10\%s^{-1}$ |
| --- | --- | --- |
| **Holzapfel et al. (2000)** | $c_1 = 0.21 \text{ kPa}; k_1^{(1)} = 122.4 \text{ kPa};$<br>$k_1^{(2)} = 59.62 \text{ kPa}; k_2 = 15.32$ | $c_1 = 64.91 \text{ kPa}; k_1^{(1)} = 205.5 \text{ kPa};$<br>$k_1^{(2)} = 12.08 \text{ kPa}; k_2 = 15.32$ |

### 4.2. Evaluation of behaviour

This study provides a unique investigation into the direction-dependent mechanical properties of the muscular layer of the human oesophagus, allowing for characterisation of the tissue’s layer-dependent and direction-dependent properties which, as of yet, have only been investigated using animal tissue.

Overall, the muscularis propria layer of the human oesophagus displayed hyperelastic properties, with greater non-linearity in the longitudinal direction than the circumferential direction. The layer was found to be anisotropic, with greater stiffness in longitudinal direction compared to the circumferential one for both strain rates used in this study. These findings are in line with those found in comparable animal studies performed by various researchers, e.g., Sommer et al. [12], Yang et al. [39] and Stavropoulou et al. [19]. They found that the longitudinal direction of both layers displayed more resistance than the circumferential direction. This can be attributed to the higher collagen content found in the longitudinal direction, as outlined in Section 3.1. The strain rate at which the tests were performed was found to affect the results in the longitudinal direction but not in the circumferential direction. This suggested strain-rate dependency is also thought to be contributed to by the increased collagen content in the longitudinal direction, as collagen fibres are often associated with the viscoelastic behaviour of soft tissues [40, 41]. The increased stiffness seen in the longitudinal direction at the higher strain rate was successfully captured by the *c*_1_ and *k*_1_ parameters of the anisotropic, hyperelastic model. The *k*_1_ parameter represents the stiffness of the anisotropic material components of the tissue, i.e. the collagen fibre families. The increase of the *k*_1_ parameter for the 10%*s*^−1^ simulation compared to 1%*s*^−1^ simulation further supports the notion that the collagen fibres contribute to the strain-rate dependency seen in the longitudinal direction. The increase of the *c*_1_ parameter at the higher strain rate suggests a non-purely elastic isotropic matrix, implying viscoelastic behaviour of this component also. Future work is needed to fully quantify any strain-rate dependency of the human oesophagus, as well as more work in investigating its viscoelastic properties.

Non-linearity of oesophageal tissue has been related to its physiological function, wherein the wall displays compliance at low strains to accommodate for the swallowing process, but stiffens at high strains in order to prevent over-dilatation [42]. This provides explanation for the non-linearity and lower stiffness seen in the circumferential direction but fails to consider the material behaviour observed in the longitudinal direction. The role of the longitudinal muscle fibres in the oesophagus have been linked to the reduction of force exerted by the circular muscle fibres required to transport the fluid bolus during peristalsis [43]. Contraction of the longitudinal muscle fibres causes the oesophagus to shorten. The stiffness of the longitudinal direction under tension, therefore, is thought to resist over-extension in the longitudinal direction during passage of the bolus so as to allow the longitudinal muscle fibres to effectively carry out their primary role of local shortening. Local shortening increases the thickness of the oesophageal wall and reduces the work of the circular muscle fibres when contracting to transport the bolus [43].

### 4.3. Considerations

Variation when testing on human soft tissues is prevalent with many variables potentially affecting their mechanical properties and therefore the mechanical data collected. These include, but are not exclusive to, age, sex, and health of the patient, as well as experimental variables such as time since explantation, sample thickness due to variation in biological composition throughout the tissue, temperature and moisture of the samples, and storage technique. In regards to the current study, variability in terms of fracture stretches and stiffness were observed between different trials. Some of these differences may be attributed to the variations seen in sample dimensions. All samples were obtained from a singular oesophagus, however, further measures were taken in order to reduce discrepancies. For instance, by obtaining samples from the inferior section of the thoracic region wherein the longitudinal muscle fibres are more evenly distributed (as outlined in Section 2.1), reducing the effect of regional differences. However, natural variation was still present, leading to the range of sample thicknesses seen in Table 2. The differences amongst the width of the samples are attributed to the human error associated with cutting samples by hand. Human error was also potentially present in measuring the width and thickness of the samples. To reduce the risk of this, measurements were taken by a singular person, and to reduce any effect, three separate measurements were taken per dimension along the sample and an average was used. Due to time constraints of the experimental setup, variations also existed in terms of the time lapsed between dissection and testing, which could further contribute to the differences seen in stress and fracture stretches of the different samples.

Some further considerations, in terms of results presented, include that the human tissue obtained for testing was extracted from an embalmed cadaver, rather than a fresh cadaver, due to the restrictions placed on postmortem testing during the COVID-19 pandemic. The cadaver was preserved in formalin after death which could affect the results, as embalming has been found to influence the stiffness of soft tissues when compared to their fresh or frozen counterparts [44, 45]. Embalming is considered to most notably affect the collagen of soft tissues, wherein the process partially denatures the fibres, causing an overall softening of the material [44]. However, formalin solution has also been found to cause collagen cross-links to form [46], which may cause an increase to the overall tissue stiffness. Girard et al. [44] found that embalming decreased the mechanical properties of the human bile duct by 80% in the longitudinal direction (more collagen-dense direction) and 40% in the circumferential direction compared to the fresh equivalents, while Hohmann et al. [45] found the Young’s modulus of the human upper biceps tendon to be approximately 20 times higher for the embalmed tendon compared to the fresh. The effect of embalming as a preservation process within literature has been found to cause mechanical properties that differ from their fresh counterparts, however it is still highly inconclusive in what way. Further to this, during layer separation, some of the collagen and elastin dense connective tissue of the submucosa may remain present on the muscular layer, potentially affecting the displayed mechanical properties. However, extreme care was taken while separating the layers to reduce the likelihood of this. Finally, the tests were conducted in air at ambient temperature rather than in a thermostatic tank. However, due to the speed of the tests conducted, the effect of a more physiological environment would have been minimal.

## 5. Conclusion

This study provides experimental data, in regards to its mechanical behaviour, on the thoracic region of the muscularis propria of the human oesophagus, in both the longitudinal and circumferential directions, and at two different strain rates. The experiments revealed hyperelastic behaviour, with greater non-linearity in the longitudinal direction; anisotropy, with greater stiffness in the longitudinal direction; and a possible strain-rate dependency of the longitudinal direction, which has been attributed to the viscoelasticity of soft tissues. The anisotropic, hyperelastic matrix-fibre model produced a good fit with the 1%*s*^−1^ experimental findings, while the parameters associated with the stiffness of the anisotropic components and the isotropic matrix could be changed to capture the difference in stiffness seen at the higher strain rate. The experimental challenges when testing human soft tissues are considered, and future work into the viscoelastic properties of the organ is suggested. The mechanical data presented has applications in tissue engineering, medical device design, as well as surgical simulations.

## Acknowledgements

C. Durcan and M. Hossain are indebted to the Swansea University Strategic Partnerships Research Scholarships (SUSPRS) for funding of the project.

